# α-Synuclein pathology disrupts mitochondrial function in dopaminergic and cholinergic neurons at-risk in Parkinson’s disease

**DOI:** 10.1101/2023.12.11.571045

**Authors:** Fanni F. Geibl, Martin T. Henrich, Zhong Xie, Enrico Zampese, Tatiana Tkatch, David L. Wokosin, Elena Nasiri, Constantin A. Grotmann, Valina L. Dawson, Ted M. Dawson, Navdeep S. Chandel, Wolfgang H. Oertel, D. James Surmeier

**Affiliations:** Department of Neuroscience, Feinberg School of Medicine, Northwestern University, Chicago, IL 60611, USA; Department of Neurology, Philipps University Marburg, Marburg 35043, Germany; Department of Psychiatry and Psychotherapy, Philipps University Marburg, Marburg 35043, Germany; Neuroregeneration and Stem Cell Programs, Institute for Cell Engineering, Johns Hopkins University School of Medicine, Baltimore, MD 21205, USA; Department of Neurology, Johns Hopkins University School of Medicine, Baltimore, MD 21205, USA; Solomon H. Snyder Department of Neuroscience, Johns Hopkins University School of Medicine, Baltimore, MD 21205, USA; Department of Physiology, Johns Hopkins University School of Medicine, Baltimore, MD 21205, USA; Department of Pharmacology and Molecular Sciences, Johns Hopkins University School of Medicine, Baltimore, MD 21205, USA; Department of Medicine, Feinberg School of Medicine, Northwestern University, Chicago, IL 60611, USA; Aligning Science Across Parkinson’s (ASAP) Collaborative Research Network, Chevy Chase, MD 20815 US

**Keywords:** Parkinson’s disease, alpha-synuclein, mitochondria, substantia nigra, dopaminergic, pedunculopontine nucleus, cholinergic, Lewy pathology, transcriptome, electrophysiology, bioenergetics, ATP

## Abstract

**Background:** Pathological accumulation of aggregated α-synuclein (aSYN) is a common feature of Parkinson’s disease (PD). However, the mechanisms by which intracellular aSYN pathology contributes to dysfunction and degeneration of neurons in the brain are still unclear. A potentially relevant target of aSYN is the mitochondrion. To test this hypothesis, genetic and physiological methods were used to monitor mitochondrial function in substantia nigra pars compacta (SNc) dopaminergic and pedunculopontine nucleus (PPN) cholinergic neurons after stereotaxic injection of aSYN pre-formed fibrils (PFFs) into the mouse brain.

**Methods:** aSYN PPFs were stereotaxically injected into the SNc or PPN of mice. Twelve weeks later, mice were studied using a combination of approaches, including immunocytochemical analysis, cell-type specific transcriptomic profiling, electron microscopy, electrophysiology and two-photon-laser-scanning microscopy of genetically encoded sensors for bioenergetic and redox status.

**Results:** In addition to inducing a significant neuronal loss, SNc injection of PFFs induced the formation of intracellular, phosphorylated aSYN aggregates selectively in dopaminergic neurons. In these neurons, PFF-exposure decreased mitochondrial gene expression, reduced the number of mitochondria, increased oxidant stress, and profoundly disrupted mitochondrial adenosine triphosphate production. Consistent with an aSYN-induced bioenergetic deficit, the autonomous spiking of dopaminergic neurons slowed or stopped. PFFs also up-regulated lysosomal gene expression and increased lysosomal abundance, leading to the formation of Lewy-like inclusions. Similar changes were observed in PPN cholinergic neurons following aSYN PFF exposure.

**Conclusions:** Taken together, our findings suggest that disruption of mitochondrial function, and the subsequent bioenergetic deficit, is a proximal step in the cascade of events induced by aSYN pathology leading to dysfunction and degeneration of neurons at-risk in PD.

## Background

PD is the second most common neurodegenerative disease, afflicting millions of people worldwide. Clinically, patients manifest a combination of motor deficits, including bradykinesia, resting tremor, and/or rigidity. These symptoms are largely attributable to the degeneration of SNc dopaminergic neurons and the resulting dysregulation of motor circuits [1]. Although symptomatic treatments are available, there are no proven strategies for stopping or slowing disease progression, reflecting an incomplete grasp of PD pathogenesis.

Intraneuronal accumulation of misfolded aSYN into proteinaceous aggregates, termed Lewy pathology (LP) is a key neuropathological feature of many types of PD [2–4]. Fibrillar forms of aSYN are thought to propagate through neuronal networks, leading to LP formation in vulnerable populations [5, 6]. However, the sequence of events set in motion by internalizing misfolded aSYN is poorly defined. The intracellular ’interactome’ of aSYN encompasses proteins and lipids involved in many aspects of cellular function, including DNA repair, vesicular trafficking, lysosomal function, and mitochondrial metabolism [7–9].

Mitochondria are particularly intriguing members of the putative aSYN interactome because of the evidence linking them to PD pathogenesis [10]. Mitochondria in at-risk neurons have elevated oxidant stress [11, 12]. Indeed, post-mortem studies of SNc dopaminergic neurons in PD patients have revealed signs of oxidant damage to mitochondrial complex I (MCI) and mitochondrial DNA [13, 14]. Loss of MCI function alone in dopaminergic neurons is sufficient to induce a progressive parkinsonian phenotype in mice [15] and environmental toxins that disrupt MCI function have been linked to PD risk [16]. Consistent with the notion that mitochondrial damage is part of the PD etiology, loss-of-function mutations in the genes coding for PINK1 (PARK6) and Parkin (PARK2), which play important roles in mitochondrial quality control, cause early-onset PD [17, 18].

There appear to be several potential sites of interaction between aSYN-induced pathology and mitochondria, ranging from proteins at the outer mitochondrial membrane and points of contact with the endoplasmic reticulum [19, 20], to inner mitochondrial membrane complexes [21–25].

However, most of these potential sites of interaction have been identified *in vitro* with cell lines or immature neurons using high concentrations of aSYN, raising concerns about its relevance to propagated pathology in an aging brain. Moreover, little of this work has focused on the types of neuron known to be selectively vulnerable to aSYN pathology in PD. This shortcoming is important because cell-type specific variation in basal stress and post-translational modifications of misfolded aSYN may be critical contributors to pathogenesis [9, 26, 27].

In an attempt to address these shortcomings, aSYN PFFs were stereotaxically injected into two PD-relevant brain regions of mice – the SNc and the pedunculopontine nucleus (PPN) – and then the consequences on mitochondria assessed using a combination of optical, transcriptomic, electrophysiological, and electron microscopic methods [11, 28]. These studies revealed that extracellular deposition of aSYN PFFs induced the formation of intracellular aggregates containing p-aSYN selectively in SNc dopaminergic and PPN cholinergic neurons. This pathology disrupted mitochondrial oxidative phosphorylation, resulting in impaired mitochondrial adenosine triphosphate (ATP) generation, elevated oxidant stress, and slowing of autonomous pacemaking. In SNc dopaminergic neurons, synucleinopathy was paralleled by a complex epigenetic reprogramming, including a broad down-regulation of mitochondrial genes. In addition, aSYN PFFs induced intracellular accumulation of large end-stage lysosomes, indicative of disturbed lysosomal function.

Together, these observations suggest that mitochondrial dysfunction is a proximal step in aSYN- induced neurodegeneration, pointing to a mechanistic convergence in PD pathogenesis.

## Methods

### Animals

All animal experiments were performed according to the NIH Guide for the Care and Use of Laboratory Animals and approved by the Northwestern University Animal Care and Use Committee. Mice were housed in groups of up to 5 animals per cage with food and water provided ad libitum on a 12-hour light/dark cycle. Throughout the study we used heterozygous DAT-Cre mice (B6.SJL- Slc6a3tm1.1(cre)Bkmn/J; Stock JAX:006660, Jackson Laboratory, USA) or heterozygous ChAT-Cre mice (B6;129S6-Chattm2(cre)lowl/J; Stock JAX:006410, Jackson Laboratory, USA), except for the electrophysiological studies for which DAT-Cre or ChAT-Cre mice were crossed in-house with Ai14- tdTomato mice (B6;129S6-Gt(ROSA)26Sortm14(CAG-tdTomato)Hze/J; Stock JAX:007908, Jackson Laboratory, USA), to allow reliable cell identification. We used a similar distribution of male and female mice in all experiments. All animals were between two and three months old at the beginning of the experiments.

### Preparation of purified aSYN PFFs and aSYN monomers

Mouse full-length aSYN PFFs were prepared at Johns Hopkins University School of Medicine, Baltimore, USA as previously described [6, 29, 30].

Briefly, purified monomeric full-length mouse aSYN was stirred at 350 rpm in a glass vial with a magnetic stir bar for 7 days. Thereafter, formed aSYN aggregates were sonicated for 30 seconds at 10% amplitude with a Branson Digital Sonifier (Danbury, CT, USA). Separation of aSYN monomers and PFFs was performed with fast protein liquid chromatography using a Superose 6 10/300GL column (GE Healthcare, Life Sciences, Wauwatosa, WI, USA). PFFs were briefly sonicated, and monomeric aSYN and PFFs were frozen at -80°C. A subset of the stock solutions was then used to perform several quality control experiments, including structure analysis with transmission electron microscopy (TEM), immunoblots, and thioflavin T (ThT) binding assays. For TEM analysis, aSYN PFFs were first adsorbed to copper grids (Electron Microscopy Sciences, Hatfield, PA, USA). After washing three times, the grids were negatively stained with uranyl formate. Images were acquired using Philips/FEI BioTwin CM120 Transmission Electron Microscope (Hillsboro, OR, USA). TEM images confirmed the fibrillar morphology of PFFs (∼100 nm). For ThT analysis, aSYN monomers and PFFs were incubated with 50 µM ThT, and fluorescence was measured at 450 nm of excitation and 485 nm of emission wavelengths by a plate reader (Tecan, Switzerland). PFFs exhibited a significantly increased fluorescence intensity compared to aSYN monomers. The ability to induce cellular aSYN pathology was further investigated by treating primary cortical neurons from C57BL/6 mice with aSYN PFFs. After 7 days of incubation, all neurons were fixed with 4% paraformaldehyde (PFA) and permeabilized with 0.2% Triton X-100. Then, cells were blocked and incubated with primary anti-aSYN (pS129) antibody (ab51253, Abcam, Cambridge, MA, USA), confirming the induction of S129 phosphorylated aSYN pathology in primary neurons. Last, aliquoted stock solutions were shipped to Northwestern University, Chicago, USA on dry ice and stored at −80°C. At Northwestern University, PFFs were thawed, and sterile phosphate-buffered saline (PBS) was added to the solution to achieve a final protein concentration of 2.5 µg/µL. Thereafter, PFFs were sonicated for 90 seconds at 10% amplitude, aliquoted and finally stored at −80 °C. On the day of injection, aliquoted PFFs were thawed and briefly vortexed before injection. **Protocol:** dx.doi.org**/10.17504/protocols.io.dm6gpbw28lzp/v1**

### Stereotaxic injections

All stereotaxic injections were conducted with a computer guided system (Angle Two, Leica Biosystems, USA) as previously described [6]. First, isoflurane-anesthetized mice were placed in the stereotaxic frame (David Kopf Instruments, USA). A total volume of 550 nl of PFFs or monomeric aSYN with a concentration of 2.5 µg/µl and 550 nl of the respective viral constructs was divided and injected in two injection spots (for SNc: AP: -3.10, ML: -1.30, DV: -4.45, and AP: - 3.50, ML: -1.30, DV: -4.20; For PPN: AP: -4.36, ML: -1.12, DV: -3.70 and AP: -4.55, ML: -1.19, DV: -3.55; for STR: AP: +0.14, ML: -2.00, DV: -2.65; AP: +0.14, ML: -2.00, DV: -3.00; AP: +0.50, ML: -1.80, DV: -2.65; AP: +0.50, ML: -1.80, DV: -3.00). To inject, a glass pipette (P-97 Pipette Puller, Sutter Instruments, Novato, CA), containing the respective viral vectors or aSYN proteins, was navigated to the injection site. After drilling a small hole, the glass pipette was slowly lowered into the brain.

Injections were performed at low speed (100 nl/min) with an automated microinjector (IM-300, Narishige, Japan), and the pipette was left for an additional 5 min in the brain after the injection was completed. Protocol: dx.doi.org/10.17504/protocols.io.81wgby191vpk/v1

### *Ex vivo* brain slice preparation

Mice were anesthetized with a mixture of ketamine (50 mg/kg) and xylazine (4.5 mg/kg) and sacrificed by transcardial perfusion with ice cold, oxygenated modified artificial cerebrospinal fluid (aCSF) containing in mM: 125 sucrose, 2.5 KCl, 1.25 NaH2PO4, 25 mM NaHCO3, 0.5 mM CaCl2, 10 mM MgCl2, and 25 mM glucose. Once perfused, the brain was rapidly removed and coronal or parasagittal slices containing the SNc or PPN region were cut using a vibratome (VT1200S Leica Microsystems). For electrophysiological experiments, coronal SNc and parasagittal PPN slices were cut 225 μm thick, whereas for two photon laser scanning microscopy (2PLSM) experiments coronal SNc and PPN slices were cut 275 μm thick, respectively. Thereafter, brain slices were incubated in oxygenated modified aCSF containing: 135.75 mM NaCl, 2.5 mM KCl, 25 mM NaHCO3, 1.25 mM NaH2PO4, 2 mM CaCl2, 1 mM MgCl2, and 3.5 mM glucose; at 34°C for 30 min, then at room temperature for another 30 min before experiments. All solutions were pH 7.4, 310–320 mOsm L^−1^ and continuously bubbled with 95% O2/5% CO2. Experiments were performed at 32–34 °C. Protocol: dx.doi.org/10.17504/protocols.io.x54v9p1e4g3e/v1

### 2PLSM imaging

Fluorescence was measured using an Ultima Laser Scanning Microscope System (Prairie Technologies) with a DODT contrast detector to provide bright field transmission images, with an Olympus 60X/1.0 NA water-dipping objective lens. A Chameleon Ultra series tunable (690- 1040mm) Ti: sapphire laser system (Coherent laser group) provided the 2P excitation source. Laser power attenuation was achieved with two Pockels’ cell electro-optic modulators (M350-80-02-BK and M350-50-02-BK, Con Optics) in series controlled by PrairieView v5.3–5.5. Non-de-scanned emission photons were detected with GaAsP photomultiplier tube (PMT) (green, 490 nm to 560 nm) and multi-alkali PMT (red, 585 nm to 630 nm).

### 2PLSM ex vivo ATP/ADP measurements

Heterozygous DAT-Cre or ChAT-Cre mice were injected with either aSYN PPFs or aSYN monomer protein as described above. 10 weeks after the initial stereotactic surgery, those mice, and another previously not injected age-matched cohort of mice (wild-type control), were injected with either 400 nl (SNc) or 250 nl (PPN) AAV9-EF1α-DIO-GW1- PercevalHR-WPRE (titer: 2.02 x 10^13^ vg/ml, from Virovek) into the SNc or PPN to induce expression of the genetically encoded ATP/ADP sensor PercevalHR. 14 days after AAV injection, mice were sacrificed and *ex vivo* brain slices were prepared as described above. Measurements were conducted as previously described [15]. Briefly, fluorescence was measured using 2PLSM as described above. To estimate the ATP/ADP ratio using PercevalHR, the probe was excited with 950 nm and 820 nm light in rapid succession. Green channel (490–560 nm) fluorescent emission signals for both wavelengths were detected using a non-descanned Hamamatsu H7422P-40 select GaAsP PMT. Two time series of 5 frames (rate of 3–4 f.p.s., 0.195 × 0.195 mm pixels and 12 µs px^−1^ dwell time) were acquired for each wavelength. Time series were analyzed offline using FIJI. The cytosol and a background region-of-interest were measured, the background was subtracted, and the 950/820 ratio was calculated for the cytosol at each time point. The contribution of mitochondria to the bioenergetic status of each cell (the OXPHOS index) was estimated by comparing the drop in the PercevalHR ATP/ADP ratio induced by bath application of oligomycin (10 μM) with the drop in the ratio after superfusion with a modified aCSF with oligomycin (10 μM) and glucose replaced with the non-hydrolyzable 2-deoxy- glucose (3.5 mM). **Protocol:** dx.doi.org/10.17504/protocols.io.8epv5xe44g1b/v1

### 2PLSM ex vivo redox measurements

Oxidant stress was assessed using a redox-sensitive roGFP probe targeted to the cytosol or the mitochondrial matrix, as previously described [31]. Therefore, heterozygous DAT-Cre or ChAT-Cre mice were injected with either aSYN PPFs or aSYN monomer protein as described above. 10 weeks after the initial stereotactic surgery, those mice, and another previously not injected age-matched cohort of mice (wild-type control), were injected with either 400 nl (SNc) or 250 nl (PPN) AAV9-CMV-DIO-rev-MTS-roGFP-WPRE (titer: 2.26 x 10^13^ vg/ml, from Virovek) for expression of mitochondria-targeted roGFP, or AAV9-CMV-DIO-rev-roGFP-WPRE (titer: 2.23 x 10^13^ vg/ml, from Virovek) for expression of cytosol targeted roGFP. 14 days after AAV injection, mice were sacrificed and *ex vivo* brain slices were prepared as described above. Slices were transferred to a recording chamber and continuously perfused with modified aCSF at 32–34 °C. Fluorescence was measured using an Ultima Laser Scanning Microscope system (Bruker) with a DODT contrast detector to provide bright-field transmission images with an Olympus ×60/0.9 NA lens. A 2P laser (Chameleon Ultra II, Coherent) tuned to 920 nm was used to excite roGFP. Time series images of the roGFP probe were acquired with 30 frames obtained over ∼20 s, with 0.197 μm × 0.197 μm pixels and 10 μs dwell time. The dynamic range of the probe was determined with 2 mM dithiothreitol, a reducing agent, and 200 μM aldrithiol, an oxidizing agent, which were used to sequentially perfuse slices. Time series images were acquired with each to determine the maximal and minimal fluorescence intensity. Time series images were analyzed offline, and fluorescence measurements in multiple regions of interest were evaluated with the background subtracted. Protocol: dx.doi.org/10.17504/protocols.io.j8nlkomb1v5r/v1

### 2PLSM ex vivo measurements of free mitochondrial Ca2+ levels

Mitochondrial free Ca^2+^ levels were measured with the mitochondrially targeted Ca^2+^-sensitive probe mito-GCaMP6. Heterozygous DAT-Cre or ChAT-Cre mice were injected with either aSYN PPFs or aSYN monomer protein as described above. 10 weeks after the initial stereotactic surgery, those mice, and another previously not injected age-matched cohort of mice (wild-type control), were injected with either 400 nl (SNc) or 250 nl (PPN) AAV9-CMV-DIO-2MT-GCaMP6 (titer: 2.22 x 10^13^ vg/ml, from Virovek) for expression of mitochondrially targeted GCaMP6. 14 days after AAV injection, mice were sacrificed and *ex vivo* brain slices were prepared as described above. Slices were transferred to a recording chamber and continuously perfused with modified aCSF at 32–34 °C. Fluorescence was measured using an Ultima Laser Scanning Microscope system (Bruker) with a DODT contrast detector to provide bright-field transmission images with an Olympus ×60/0.9 NA lens. A 2P laser (Chameleon Ultra II, Coherent) tuned to 920 nm was used to excite mito-GCaMP6. Time series images of the GCaMP6 probe were acquired with 30 frames obtained over ∼20 s, with 0.197 μm × 0.197 μm pixels and 10 μs dwell time. The dynamic range of the probe was determined as follows. First, sections were perfused with a modified aCSF in which Ca^2+^ was substituted with Mg^2+^ (0 mM Ca^2+^, 3mM Mg^2+^) and the Ca^2+^ ionophore ionomycin was added at a concentration of 1 µM. Hereafter, a modified aCSF containing 3 mM Ca^2+^ and 1µM ionomycin was perfused. Time series images were acquired with each to determine the minimal and maximal fluorescence intensity. Time series images were analyzed offline, and fluorescence measurements in multiple regions of interest were evaluated with the background subtracted. **Protocol:** dx.doi.org**/10.17504/protocols.io.5jyl8pby7g2w/v1 Electrophysiology:** For electrophysiological experiments, DAT-Cre-Ai14-tdTomato and ChAT-Cre- Ai14-tdTomato mice were used to allow reliable identification of SNc dopaminergic or PPN cholinergic neurons. Conventional tight-seal (>2 GΩ) patch-clamp recordings were made in cell-attached mode from either aSYN PFF or aSYN monomer injected mice, or age-matched non-injected (wild-type) mice. *Ex vivo* brain slices were prepared as described above. After slicing, 225 µm thick sections were stored in a holding chamber at room temperature filled with modified aCSF and continuously bubbled with 95% O2 and 5% CO2. Patch pipettes were pulled from thick-walled borosilicate glass pipettes on a P-1000 puller (Sutter Instrument). Pipette resistance was typically 4-5 MΩ. Recording pipettes were filled with modified aCSF. Synaptic blockers were bath-applied for all recordings (10 µM DNQX, 50 µM D-AP5, 10 µM SR 95531 hydrobromide, 1 µM CGP55845 hydrochloride). Recordings were obtained at 32-34 °C temperature, using a MultiClamp 700A amplifier (Molecular Device) and pClamp 10 software (Molecular Device). Data were acquired at 100 kHz, filtered at 10 kHz, digitized using a DigiData 1440 (Molecular Devices), and analyzed using Clampfit 10.3 (Molecular Devices). **Proto**col: dx.doi.org/10.17504/proto**cols.io.eq2lyj8zrlx9/v1 RiboTag profiling and RNAseq:** Heterozygous DAT-Cre mice were injected with either aSYN PFFs or aSYN monomer protein into the SNc as described above. 10 weeks after the initial injection, all mice were injected with 400 nl of AAV5- EF1a-DIO-Rpl22ll-3xFlag-2A-GFP-WPRE (Virovek; titre of 2.13 x 10^13^ vg/ml) into the SNc. Four weeks after the injection, mice were anesthetized with a mixture of ketamine (50mg/kg) and xylazine (4.5mg/kg) and sacrificed by transcardial perfusion with ice cold aCSF. Brains were quickly removed from the skull and 225 μm thick coronal slices containing the SNc region were cut using a vibratome (VT1200S Leica Microsystems). Thereafter, the SNc region was dissected from the tissue slices and immediately frozen at −80 °C. RiboTag immunoprecipitation was performed as described previously [15]. In brief, tissue was homogenized in cold homogenization buffer (50 mM Tris (pH 7.4), 100 mM KCl, 10 mM MgCl2, 1 mM dithiothreitol, 100 μg ml^−1^ cycloheximide, protease inhibitors and recombinant RNase inhibitors, and 1% NP-40). Homogenates were centrifuged 10,000g for 10 min, and the resulting supernatant was precleared with protein G magnetic beads (Thermo Fisher Scientific) for 1 h at 4 °C with constant rotation.

Immunoprecipitations were carried out with anti-Flag magnetic beads (Sigma-Aldrich) at 4 °C overnight with constant rotation. Four washes were carried out with high-salt buffer (50 mM Tris (pH 7.4), 350 mM KCl, 10 mM MgCl2, 1% NP-40, 1 mM dithiothreitol and 100 μg ml^−1^ cycloheximide). RNA extraction was performed using the RNA-easy Micro RNA extraction kit (QIAGEN) according to the manufacturer instructions. Protocol: dx.doi.org/10.1**7504/protocols.io.261gedwyyv47/v1**

### RNASeq analysis

RNASeq analysis was performed as previously described [15]. The quality of reads, in FASTQ format, was evaluated using FastQC. Reads were trimmed to remove Illumina adapters from the 3′ ends using cutadapt. Trimmed reads were aligned to the *mus musculus* genome (mm10) using STAR [32]. Read counts for each gene were calculated using htseq-count [33] in conjunction with a gene annotation file for mm10 obtained from Ensembl(http://useast.ensembl.org/index.html). Normalization and differential expression were calculated using DESeq2, which uses the Wald test [34]. The cut-off for determining significantly differentially expressed genes was a false-discovery-rate-adjusted P value of less than 0.05 using the Benjamini–Hochberg method. To reduce noise from contamination via glial and non-dopaminergic neuronal ribosomes, analysis was limited to genes expressed at least 200 counts. For pathway analysis, GSEA software (https://www.gsea-msigdb.org/gsea/index.jsp) [35] was used. Heatmaps were generated using Morpheus software (https://software.broadinstitute.org/morpheus).

### Histology and imaging

All tissue processing and immunohistochemistry was performed as previously described [6]. To achieve a sufficient fixation of the brain for further immunohistochemical analysis, all mice were anesthetized with a mixture of ketamine (50mg/kg) and xylazine (4.5mg/kg) and sacrificed by transcardial perfusion with ice cold 0.1 M PBS followed by 4% ice-cold PFA for 5 min.

After perfusion, mice were decapitated and brains were quickly removed, followed by post-fixation for 3 days in PFA and 3 days in 30% sucrose solution in 0,1 M PBS. Brains were then frozen on dry ice and stored in -80°C until sectioning. On the day of sectioning, brains were embedded in tissue freezing media (OCT Compound, Tissue Tek, USA) and cut into 30 μm thick consecutive coronal sections using a cryostat microtome (CM3050 S, Leica, Germany). All sections spanning the complete rostro-caudal extent of the brain were kept in correct order and stored at 4°C in cryoprotect-solution (1:1:3 volume ratio of ethylenglycol, glycerol and 0.1 M PB) until further processing.

Immunofluorescence stainings used for data analysis or representative images were performed according to the following protocol. Sections were washed 4 x 5 min in 0.1 M PB buffer and blocked for 1 hour in 10% normal donkey serum (NDS) in 0.1 M PB with 0.3% Triton X-100 (PBT) at room temperature (RT). Primary antibodies were diluted in 10% NDS in PBT and incubated overnight at 4°C. On the second day, sections were washed 4 x 5 min in PBT, then incubated with fluorophore-conjugated, species-specific secondary antibodies for 2 hours at RT, blocked with 10% NDS in PBT. In most cases, sections were additionally stained with DAPI (DAPI, Sigma-Aldrich, D9542-5MG, 1:10,000 of 5 mg/ml) for 10 min in 0.1 M PB. Before mounting with antifade mounting medium (ProLong Diamond Antifade Mountant, Invitrogen, P36965), sections were washed 5 x 5 min in PBT. Exceptions to this general immunofluorescence staining protocol were made for staining of S129-phosphorylated aSYN, where a streptavidin-based amplification of fluorescence was used. For this, sections were washed 4 x 5 min in 0.1 M PB, and blocked for 1 hour in 10% NDS in PBT at RT. The primary antibody (anti-aSYN (pS129), Abcam, ab51253) was diluted in 10% NDS in PBT and incubated overnight at 4°C. On the second day, after an initial wash for 4 x 5 min in PBT, sections were incubated with a biotinylated species-specific secondary antibody (biotinylated anti-rabbit, Jackson ImmunoResearch, 711-065-152, 1:1000) directed against the rabbit p-aSYN antibody with 10% NDS in PBT for 1 hour at RT. Sections were then washed (3 x 5 min in PBT) and incubated with fluorophore-conjugated streptavidin (Streptavidin AlexaFluor647, Jackson ImmunoResearch, 016-600-084, 1:1000) in 10% NDS in PBT for 2 hours at RT. Before mounting, sections were additionally stained with DAPI as described above and thereafter washed again 5 x 5 min in PBT. Representative fluorescent images were acquired with a TCS SP8 confocal microscope (Leica, Germany). All images were processed with FIJI to enhance signal-to-noise or to rearrange colors of certain image channels. **Protocol:** dx.doi.org/10.17504/protocols.io.bp2l6xpr1lqe/v1

### Proteinase K treatment

To analyze the formation of insoluble p-aSYN aggregates, SNc and PPN sections were digested with Proteinase K (PK) using a protocol described previously [36]. Briefly, sections containing the SNc and PPN regions were washed in 0.1 M PB and subsequently digested at 65°C for 10 min in PBT and 12 µg/ml PK (Proteinase K, Invitrogen, #4333793). To visualize insoluble aggregates, digested sections were double stained against p-aSYN and p62 in combination with TH or ChAT, following the fluorescence staining protocol described above. Complete absence of TH and ChAT immunoreactivity indicated successful PK digestion. Sections in which TH or ChAT immunoreactivity was still visible were excluded from analysis due to incomplete PK digestion.

Control sections received the same treatment without incubating them in PK. Protocol: dx.doi.org/10.17504/protocols.io.81wgbx9qqlpk/v1

### Quantification of SNc and PPN neuronal cell counts

To quantify TH-positive and NeuN-positive cells in the SNc, 30 μm thick coronal sections of 5 defined Bregma coordinates were analyzed (-2.92, -3.16, -3.40, 3.64, -3.80). To assess cell counts of ChAT-positive and NeuN-positive PPN neurons, the following 5 defined Bregma coordinates were analyzed: -4.24, -4.48, -4.72, -4.84, -4.96. First, brain tissue was washed 3 x 5 min in 0.1 M PB and thereafter quenched with 3% H2O2 and 10% methanol for 15 minutes at RT. After a second wash (4 x 5 min 0.1 M PB), sections were blocked for 1 hour in 5% NDS in PBT. Primary antibody (anti neuronal nuclei (NeuN), Merck Millipore, MAB377, 1:1000) was diluted in 5% NDS in PBT and incubated overnight at 4°C. On the second day, sections were washed 4 x 5 min in 0.1 M PB then incubated with a biotinylated secondary antibody (biotinylated donkey anti-mouse, Jackson ImmunoResearch, 715-065-151, 1:500) for 1 hour at RT, followed by incubation in avidin-biotin-peroxidase solution (Vectastain Elite ABC HRP Kit, Vector Laboratories, PK- 6100) for 1 hour at RT. Color reaction was initiated with 5% DAB (3,3’-Diaminobenzidin, Serva,

Cat#18865.02), diluted in 0.1 M PB with 0.02% H2O2. After color reaction, sections were washed 4 x 5 min and blocked again for 1 hour in 5% NDS in PBT. Tissue sections were then incubated with another primary antibody (for SNc sections: anti-tyrosine hydroxylase (TH), Merck Millipore, AB152, 1:1000; for PPN sections: anti-choline acetyltransferase (ChAT), Merck Millipore, AB144P, 1:100) diluted in 5% NDS in PBT overnight at 4°C. On the third day, sections were washed 4 x 5 min in 0.1 M PB, incubated with a biotinylated secondary antibody (biotinylated donkey anti-rabbit, Jackson ImmunoResearch, 711-065-152, 1:500; or biotinylated donkey anti-goat, Jackson ImmunoResearch, 705-065-147, 1:500) for 1 hour at RT, followed by incubation in avidin-biotin-peroxidase solution (Vectastain Elite ABC HRP Kit, Vector Laboratories, PK-6100) for 1 hour at RT and initiation of color reaction with a Peroxidase Substrate Kit (SG Peroxidase Substrate Kit, Vector Laboratories, SK-4700). All stained sections were mounted, dehydrated and coverslipped with mounting medium (Eukitt medium, Sigma-Aldrich, Cat#03989). Brightfield images were acquired using an AxioImager M2 microscope (Zeiss, Germany) equipped with an Axiocam 506 color camera (Zeiss, Germany). For quantification of neuronal cell counts, the optical fractionator workflow (StereoInvestigator version 9, MicroBrightField Biosciences, USA) was used. For analysis of SNc neurons, contours were drawn based on the cytoarchitectonic distribution of TH-positive neurons, whereas for assessing PPN cell counts the distribution of ChAT-positive neurons was used. Parameters used for counting were: grid size 100 × 100 μm, counting frame 85 × 85 μm, and 2 μm guard zones. **Protocol:** dx.doi.org**/10.17504/protocols.io.e6nvwdq47lmk/v1**

### Transmission electron microscopy (TEM)

For TEM experiments, we chose a projection-based approach for reliable cell identification under the TEM [15]. Therefore, heterozygous DAT-Cre mice having received either aSYN PFFs or monomers 11 weeks prior, or age matched non-injected DAT-Cre mice, were injected with 400 nl of 1% Fluorogold (Santa Cruz) dissolved in saline at a rate of 100 nl per min into the dorsal striatum. The total volume was evenly distributed over four injection sites using the following Bregma coordinates: 1. AP: +0.14, ML: -2.00, DV: -2,65; 2. AP: +0.14, ML: -2.00, DV: -3.00; 3. AP: +0.50, ML: -1.80, DV: -2.65; 4. AP: +0.50, ML: -1.80, DV: -3.00. Five days later, mice were perfused at room temperature with 0.1 M PBS followed by a fixative containing 2% paraformaldehyde, 1.25% glutaraldehyde and 0.1 M phosphate buffer (pH 7.3). Thereafter, mice were decapitated, and brains quickly removed, followed by post-fixation for 24 hours in the same fixative. 500 µm thick slices were cut using a vibratome and slices were visualized under the fluorescence microscope to localize the SNc, containing cells labelled with retrogradely transported Flurogold. Identified SNc region was then dissected and postfixed in buffered 2% OsO4, rinsed and stained in 1% uranyl acetate, dehydrated, and embedded in EMBed 812 (EMS, 14120). Ultrathin sections (90 nm) were contrasted with lead citrate and uranyl acetate and analyzed under a JEOL 1230 transmission electron microscope operated at 80 kV with a Gatan Orius CCD camera with the

Digital Micrograph software. SNc neurons were identified by their classical morphological appearance in combination with electron dense Fluorogold particles within lysosomes [15]. The nuclei, cytoplasm, mitochondria, and lysosomes were manually outlined and analyzed using FIJI. For analysis, mitochondria were classified as healthy, swollen or degenerated based on their morphological appearance (shape, christae structure, intact double membrane). **Protocol:** dx.doi.org**/10.17504/protocols.io.q26g7pw88gwz/v1**

### Statistical methods

Data analysis was performed using GraphPad Prism (version 8.3.1 GraphPad Software, USA), Clampfit 10.3 (Molecular Devices), GSEA, or FIJI. In all experiments, sample size was based on prior studies using similar techniques. Statistical analyses were performed using GraphPad Prism. Data are presented as box-and-whisker plots depicting the median, quartiles and range.

Datasets were not assumed to be normal and were therefore analyzed using nonparametric statistics. Exact statistical tests are indicated in each figure legend. Differences were considered significant at P < 0.05. Reproducibility: all key experiments were independently reproduced by different co-authors. Each experiment was performed multiple times across multiple mice as described in the figure legends. All findings in this manuscript were replicated across animals/brain slices/cells, and data were pooled for analysis and presentation. All figures were created with Adobe Illustrator version 25.1 (Adobe Systems).

## Results

### aSYN PFFs induced seeding of Lewy pathology-like inclusions and neuronal loss in SNc

As a first step towards assessing the potential impact of pathological aSYN on mitochondrial function in vulnerable neurons, mouse PFFs (or monomeric mouse aSYN) were stereotaxically injected into the SN of mice expressing Cre recombinase (Cre) under control of the dopamine transporter (DAT) promoter (DAT-Cre) (**Fig. 1a**). Twelve weeks later, DAT-Cre mice were sacrificed, their brains fixed and sectioned for histology. At this time point, roughly 40% of tyrosine hydroxylase (TH) positive SNc dopaminergic neurons manifested immunoreactivity for aSYN phosphorylated at serine 129 (pS129) (**Fig. 1b, c**) – a commonly used marker of intracellular aSYN aggregation [37, 38]. Immunoreactivity for pS129 aSYN was distributed over the anterior-posterior axis of the SNc (**Fig. S1a**). In contrast, an equivalent striatal injection of aSYN PFFs induced a much lower level of SNc pS129 immunoreactivity (**Fig. 1c**). PFF-induced α-synucleinopathy was resistant to proteinase K (PK) digestion and immunoreactive for p62, both of which are features of human LP (**Fig. 1d**) [39, 40].

**Fig. 1.**
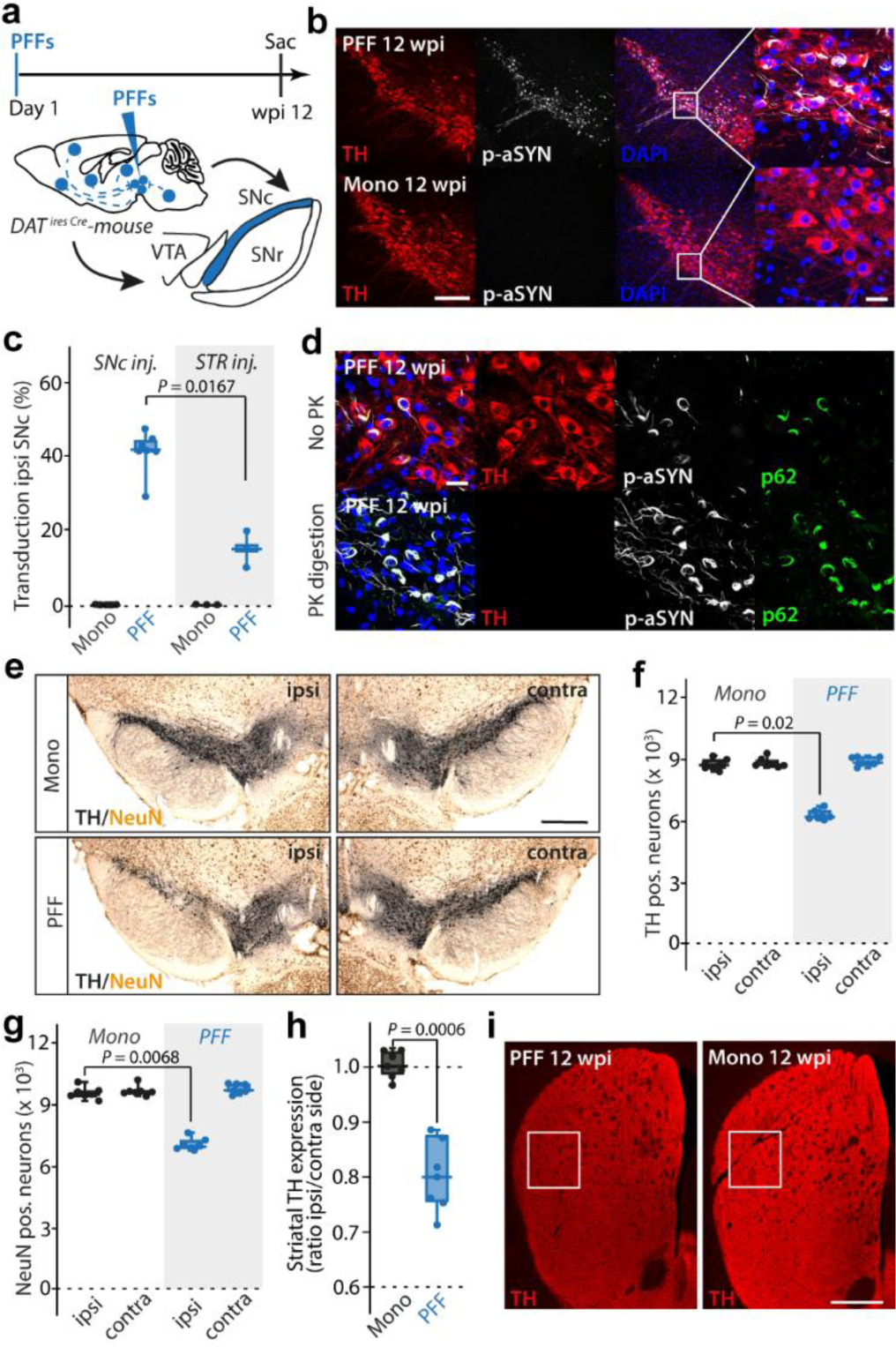
α-synuclein fibril injection causes PD-like neurodegeneration of midbrain DA SNc neurons. a, Experimental protocol. **b,** TH^+^ SNc neurons exhibiting p-aSYN pathology 12 weeks after initial seeding (upper row), while pathology is absent in *α*-synuclein monomer injected mice (lower row). Scale bar, 150 µm in overviews, 30 µm in magnified images. **c,** Transduction rates for SNc injected mice compared to conventional striatal (STR) injected mice (n = 7 for SNc, and n = 3 for STR; Mann-Whitney-U test; median *±* min/max). **d,** p- aSYN^+^ aggregates were p62^+^ and resistant to digestion with Proteinase K (PK). Scale bar, 30 µm. **e,** Representative image showing neurodegeneration of DA SNc population on the injected side. Scale bar, 500 µm. **f,** Box plot showing number of TH^+^ neurons in SNc (n = 7 for both groups; Kruskal-Wallis; median ± min/max). **g,** Box plot showing number of NeuN^+^ neurons in SNc (n = 7 for both groups; Kruskal-Wallis; median ± min/max). **h,** Quantification of TH expression in the dorsal striatum (n = 7 for both groups; Mann-Whitney-U test; median ± min/max). **i,** Image showing TH expression in striatum. White square indicates measured ROI. Scale bar, 600 µm.

Unbiased stereological quantification of TH immunoreactive (TH^+^) neurons in the SNc revealed that PPFs reduced the number of TH^+^ neurons by about a third (**Fig. 1e,f**). To determine if the reduction in TH immunoreactivity was attributable to frank cell loss or phenotypic down-regulation, the pan-neuronal marker NeuN [41] was used to estimate the number of surviving SNc neurons. The percent reduction in NeuN^+^ neurons was similar to that of TH^+^ neurons, indicating that PFFs had induced frank neurodegeneration (**Fig. 1e,g**). As expected from the somatodendritic loss, axonal TH immunoreactivity in the striatum was significantly reduced (**Fig. 1h,i**).

Despite the spread of pS129 aSYN pathology throughout the SNc and neighboring ventral tegmental area (VTA), there was no discernible pathology in neighboring non-dopaminergic neuronal populations, like those in the substantia nigra pars reticulata (SNr) or red nucleus.

### aSYN pathology disrupted mitochondrial function

Misfolded forms of aSYN have been hypothesized to disrupt cellular function in a variety of ways that might ultimately lead to neurodegeneration [9]. Although mitochondrial dysfunction figures prominently in many of these hypotheses, the ability of *in vivo* seeding of aSYN pathology to affect them has not been systematically explored. To begin filling this gap, a combination of cellular and molecular approaches was used to study surviving SNc dopaminergic neurons in *ex vivo* brain slices from mice 12 weeks after PFF injections. To assess the ability of mitochondria to generate ATP through oxidative phosphorylation (OXPHOS), SNc dopaminergic neurons were induced to express a genetically encoded, ratiometric sensor (PercevalHR) of the cytoplasmic ATP to adenosine diphosphate (ADP) ratio [15, 42]. To this end, DAT-Cre mice were stereotaxically injected with adeno-associated virus (AAV) carrying a double-floxed inverse open reading frame (DIO) expression construct for PercevalHR 10 weeks after aSYN PFF (or monomer) injection (**Fig. 2a**). Histological analysis of these mice two weeks later confirmed the colocalization of TH, PercevalHR and pS129 aSYN pathology in SNc neurons (**Fig. 2b**). In *ex vivo* brain slices, 2PLSM was used to determine the PercevalHR fluorescence (following excitation with 950 and 820 nm light) and the corresponding cytosolic ATP/ADP ratio [15]. In mice without an aSYN injection (WT) or in mice with aSYN monomer (mono) injection, inhibition of mitochondrial complex V (MCV) by bath application of oligomycin resulted in a precipitous drop in cytosolic ATP/ADP ratio (**Fig. 2c,d**), consistent with previous work showing that mitochondrial OXPHOS is central to the bioenergetic state of healthy SNc dopaminergic neurons [43]. However, in SNc dopaminergic neurons from PFF injected mice, the cytosolic ATP/ADP ratio was either unaffected or actually rose following MCV inhibition, suggesting that mitochondria had ceased making a significant contribution to cytosolic ATP regeneration. In these neurons, the cytosolic ATP/ADP ratio had become dependent upon glycolysis, as bath application of 2- deoxyglucose (2-DG) caused a profound drop in the PercevalHR ratio (**Fig. 2c,d**).

**Fig. 2.**
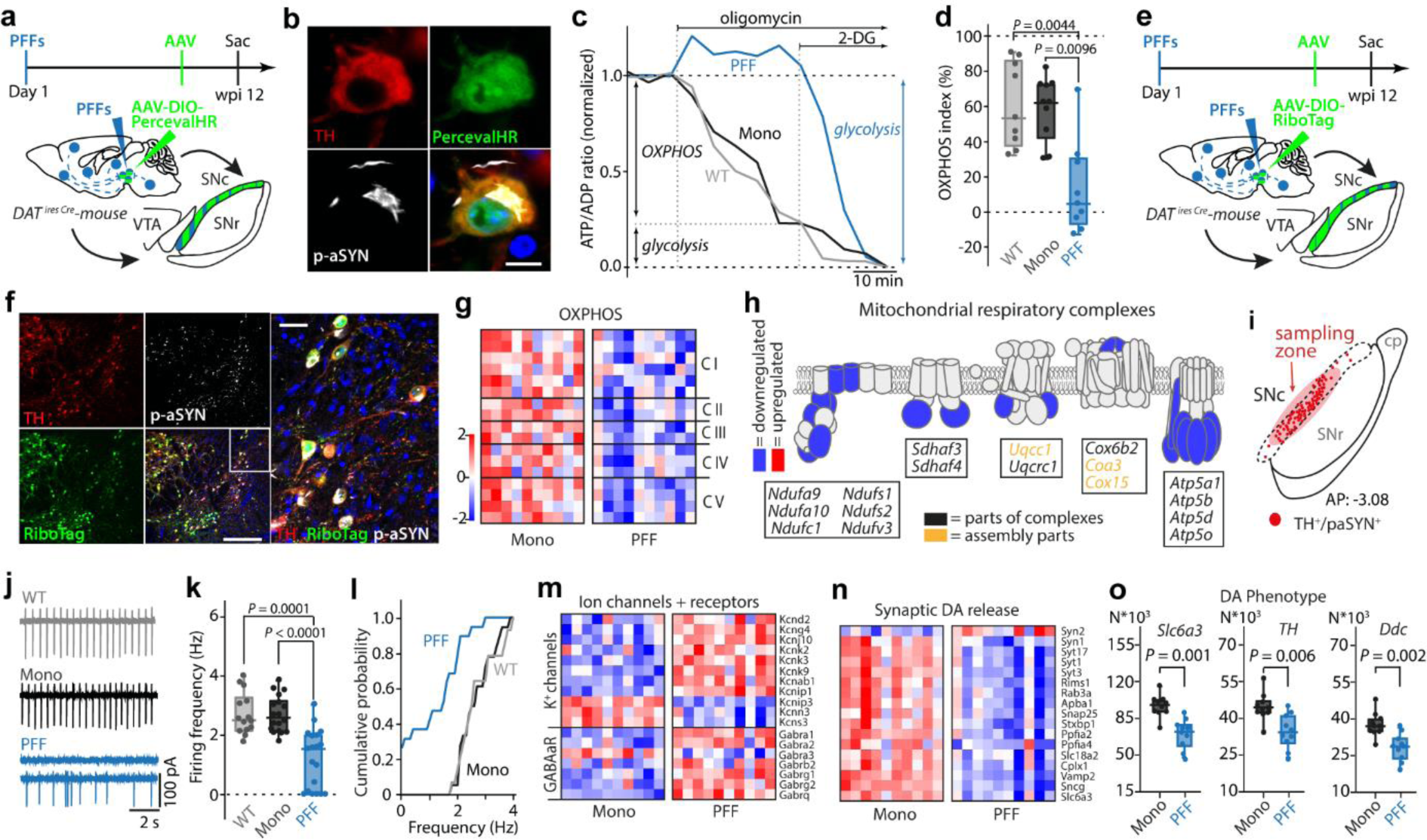
α-synucleinopathy causes disruption of mitochondrial OXPHOS resulting in energetic disbalance of DA SNc neurons. a, Experimental protocol for PercevalHR experiment. **b,** PercevalHR expression in a p-aSYN^+^ TH+ DA SNc neuron. Scale bar, 10 µm. **c,** Representative time-lapse measurements of PercevalHR fluorescence ratio. Oligomycin and 2-deoxyglucose (2-DG) were applied to determine OXPHOS and glycolytic contribution to ATP/ADP ratio. **d,** Box plot showing OXPHOS index for WT, Mono, and PFF treated mice. Synucleinopathy shifts neuronal metabolism to the glycolytic pathway while mitochondria become net consumers of ATP (n = 9/10/9 neurons from 5/6/8 mice for PFF, monomeric aSYN, and WT; Kruskal-Wallis; median ± min/max). **e,** Experimental protocol for RiboTag experiment. **f,** Image depicting expression of RiboTag in p-aSYN^+^ DA SNc neurons. Scale bar in overviews 300 µm, and 50 µm in magnified image. **g,** Heatmap of RNASeq analysis showing significantly down- or upregulated genes of OXPHOS (n = 10 for both groups; Wald test adjusted using Benjamini-Hochberg method, p < 0.05). **h,** Scheme depicting significantly down- or upregulated units of the mitochondrial respiratory chain. **i,** Scheme indicating sampling zone within SNc. **j,** Representative cell-attached recordings of DA SNc neurons. **k,** Autonomous pacemaking of DA SNc neurons (n = 19/18/14 neurons from 5/5/5 mice for PFF, monomeric aSYN, and WT; Kruskal-Wallis; median ± min/max). **l,** Cumulative probability plot of SNc DA autonomous discharge rates (n = 19/18/14 neurons from 5/5/5 mice for PFF, monomeric aSYN, and WT). **m, n,** Heatmap of RNASeq analysis showing expression profiles of K+ channel and GABAa Receptor units (m) and downregulation of synaptic DA release genes (n) (n = 10 for both groups; Wald test adjusted using Benjamini-Hochberg method, p < 0.05). **o,** Box plots showing normalized gene expression values for DA phenotype genes (n = 10 for both groups; Wald test adjusted using Benjamini-Hochberg method; median ± min/max).

To gain a better understanding of the mechanisms responsible for this metabolic shift, actively transcribed messenger ribonucleic acids (mRNAs) were harvested from SNc dopaminergic neurons in aSYN monomer or PFF injected mice and then subjected to RNASeq analysis [44]. To selectively harvest mRNAs from dopaminergic neurons, DAT-Cre mice were stereotaxically injected with an AAV vector carrying a DIO-RiboTag expression construct (**Fig. 2e**). Histological analysis of these mice confirmed the robust co-localization of TH, pS129 aSYN and the RiboTag reporter (**Fig. 2f**). Consistent with the functional studies, gene set enrichment analysis (GSEA) (**Fig. S2a,b**) revealed a broad down- regulation in the expression of genes coding for OXPHOS-related proteins in dopaminergic neurons from aSYN PFF injected mice (**Fig. 2g,h**; **Fig. S2c**). Although this feature of the translatomes of dopaminergic neurons resembled that seen following loss of MCI function by deletion of *Ndufs2* [15], other features differed. Of particular note, there was not an up-regulation of genes coding for proteins participating in glycolysis and genes associated with lactate metabolism were down- regulated (**Fig. S2c,d)**. Interestingly, genes associated with beta-oxidation of fatty acids and ketone body consumption were upregulated (**Fig. S2c,d)**. In addition to the reprogramming of metabolic pathways, GSEA revealed upregulation of apoptosis related genes, and genes linked to hypoxia (**Fig. S2e,f**). *Ex vivo* electrophysiological examination of SNc dopaminergic neurons revealed that PFF exposure suppressed autonomous pacemaking (**Fig. 2i-l**). On the transcriptomic level, these electrophysiological changes were associated with an up-regulation of genes coding for plasma membrane K^+^ channels and GABAA receptors (**Fig. 2m; Fig. S2f**) and down-regulation of genes associated with the synthesis, synaptic release and uptake of dopamine (*Slc6a3, Th, Ddc*) (**Fig. 2n,o**).

### aSYN PFFs elevated oxidant stress, lowered mitochondrial mass and induced lysosomal pathology

To assess the impact of aSYN PFF-induced pathology on mitochondrial and cytosolic redox status, genetically encoded redox-sensitive variants of green fluorescent protein (roGFP) were targeted either to the mitochondrial matrix (mito-roGFP) or the cytosol (cyto-roGFP) [31]. Ten weeks after the initial delivery of aSYN PFFs or monomers, DAT-Cre mice underwent a second stereotaxic surgery in which an AAV carrying a DIO mito-roGFP or cyto-roGFP expression construct was injected into the SNc (**Fig. 3a**). Histological analysis of these mice two weeks later confirmed the colocalization of TH, mito-roGFP and pS129 aSYN (**Fig. 3b**). 2PLSM then was used in *ex vivo* brain slices to measure mitochondrial oxidant stress (**Fig. 3c**). These experiments revealed that mitochondrial oxidant stress was elevated in SNc dopaminergic neurons following aSYN PFF exposure, but not following exposure to monomers (**Fig. 3d**). The elevation was not attributable to increased mitochondrial Ca^2+^ loading, as matrix Ca^2+^ concentration did not appear to be altered by PFFs (**Fig. S3a-c**). Consistent with previous work *in vitro* [45], aSYN PFFs also elevated cytosolic oxidant stress (measured with the cyto-roGFP probe) (**Fig. 2e,f**). On the transcriptomic level these alterations were paralleled by upregulation of *ATF5* (**Fig. S3e**), a key transcription factor of the cellular induced stress response (ISR).

**Fig. 3.**
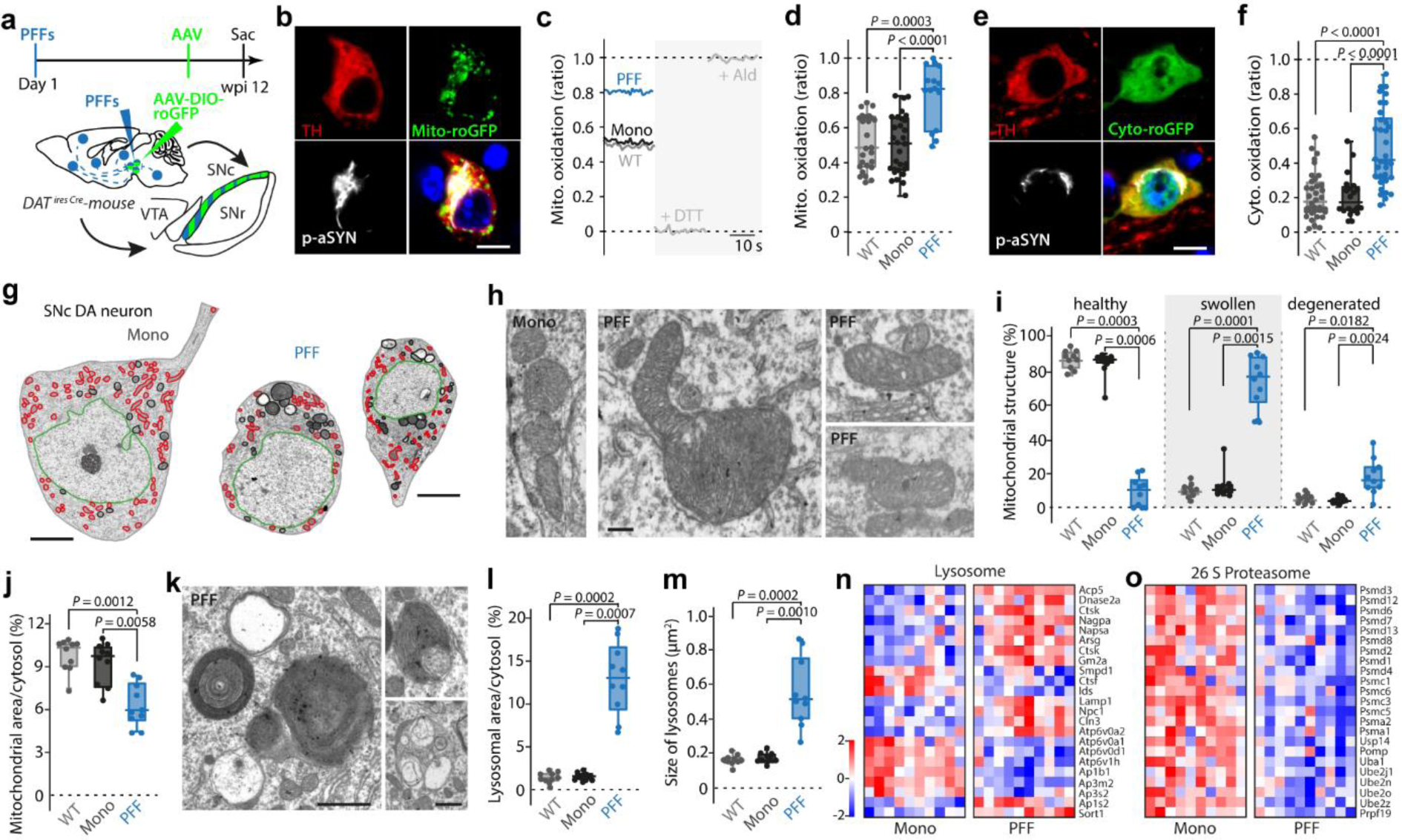
α-synucleinopathy leads to mitochondrial oxidation and morphological alterations of mitochondria and lysosomes. a, Experimental protocol. **b,** Mito-roGFP expression in a p-aSYN^+^ TH^+^ DA SNc neuron. Scale bar, 10 µm. **c,** Calibration protocol. **d,** Synucleinopathy elevates mitochondrial ROS levels (n = 15/30/26 neurons from 5/5/5 mice for PFF, monomeric aSYN, and WT; Kruskal-Wallis; median ± min/max). **e,** Cyto-roGFP expression in a p-aSYN+ TH+ DA SNc neuron. Scale bar, 10 µm. **f,** Synucleinopathy increases basal cytosolic oxidation (n = 36/20/34 neurons from 5/5/5 mice for PFF, monomeric aSYN, or WT; Kruskal-Wallis; median ± min/max). **g,** Transmission electron micrographs of SNc DA neurons from monomeric aSYN (left) or PFF (right) injected mice. The nucleus is highlighted in green, mitochondria in red, and lysosomes in black, respectively. Scale bar 10 *μ*m. **h,** Series of transmission electron micrographs showing healthy, swollen, and degenerated mitochondria of SNc DA neurons. Scale bar 300 nm. **i,** Box plots showing quantification of mitochondrial morphology (n = 10/10/10 neurons from 4/4/4 mice for PFF, monomeric aSYN, and WT; Kruskal-Wallis; median ± min/max). **j,** Box plot indicating mitochondrial density as percent of cytosol area (n = 10/10/10 neurons from 4/4/4 mice for PFF, monomeric aSYN, and WT; Kruskal-Wallis; median ± min/max). **k,** Transmission electron micrographs showing different lysosome stages, including multilamellar bodies, in a SNc DA neuron from a PFF injected mouse. Scale bar 1 *μ*m. **l,** Quantification of lysosome density as percent of cytosol area (n = 10/10/10 neurons from 4/4/4 mice for PFF, monomeric aSYN, and WT; Kruskal-Wallis; median ± min/max). **m,** Box plot indicating average size of lysosomes (n = 10/10/10 neurons from 4/4/4 mice for PFF, monomeric aSYN, and WT; Kruskal-Wallis; median ± min/max). **n, o,** Heatmaps of RNASeq analysis showing significantly down- or upregulated genes of lysosomal (n), and proteasomal (o) degradation pathways (n = 10 for both groups; Wald test adjusted using Benjamini-Hochberg method, p < 0.05).

Oxidant stress can damage proteins, lipids and deoxyribonucleic acid (DNA). To assess the structural impact of sustained oxidant stress on SNc dopaminergic neurons, they were retrogradely labeled by striatal injection of Fluorogold, and then mice were sacrificed and processed for transmission electron microscopy (TEM) (**Fig. S4a**). aSYN PFF exposure reduced the somatic cross-sectional area of SNc dopaminergic neurons (**Fig. 3g; Fig. S4b,c**). Mitochondria in PFF-exposed neurons commonly were dysmorphic and had altered cristae structure (**Fig. 3h,i; Fig. S4d)**. In addition, mitochondrial density and the total cytosolic area mitochondria occupied was significantly reduced by aSYN PFFs (**Fig. 3j; Fig. S4e,f**). SNc dopaminergic neurons from monomer or non-injected DAT-Cre-WT mice did not display any of these features.

In contrast, aSYN PFFs increased the proportion of the cytosol occupied by lysosomes, which was largely attributable to increased lysosomal size (**Fig. 3k-m, Fig. S4g,h**). The cytoplasm of PFF-exposed SNc dopaminergic neurons was crowded with endolysosomes, autophagosomes, and terminal lysosomes having a characteristic multilamellar appearance (**Fig. 3k; Fig. S4i**). Occasionally, mitochondria were found in these multilamellar bodies (**Fig. S4j**). The aSYN PFF-induced lysosomal adaptation was accompanied by a complex alteration in gene expression. Genes coding for lysosomal enzymes and structural proteins were upregulated, but genes coding for ATPases and adapter proteins were down-regulated (**Fig. 3n; Fig. S4k**). Also, genes coding for proteasomal proteins were broadly down-regulated (**Fig. 3o, Fig. S4k**), in agreement with previous reports of compromised proteasomal capacity in the presence of synuclein pathology [46, 47].

### aSYN PFFs induced similar bioenergetic changes in PPN cholinergic neurons

Previous studies have suggested that dopaminergic neurons were at elevated risk of neurodegeneration in PD because of an interaction between oxidized forms of dopamine and aSYN [12, 48]. However, aSYN pathology in the brains of PD patients is found in a variety of non-dopaminergic neurons that ultimately degenerate. For example, cholinergic neurons in the brainstem PPN not only manifest LP in PD but are lost in the course of the disease, much like SNc dopaminergic neurons [28, 49]. To determine if seeded α-synucleinopathy induced a similar set of bioenergetic alterations in these neurons, aSYN PFFs were injected into the PPN of mice expressing Cre recombinase under the control of the promoter for choline acetyltransferase (ChAT) to allow cell-specific measurements to be performed (ChAT-Cre) (**Fig. S5a**). Twelve weeks after seeding, the majority of PPN cholinergic neurons manifested p-aSYN pathology that was PK resistant (**Fig. S5b,g**). Unbiased stereological counts of ChAT-immunoreactive and NeuN^+^ PPN neurons confirmed that aSYN PFF exposure induced significant neurodegeneration (**Fig. S5d-f**). Using the same strategy described above employing genetically encoded biosensors to assess mitochondrial function in surviving PPN cholinergic neurons (**Fig. 4a,b**), we found that aSYN PFF exposure compromised the ability of mitochondrial OXPHOS to generate ATP (**Fig. 4c,d**), which was accompanied by comparable alterations of PPN cholinergic neurons electrophysiological properties, including slowing of pacemaking and an increase in irregularity of discharge (**Fig. 4e-g**). As in dopaminergic SNc neurons, pathological aSYN caused increased levels of mitochondrial and cytosolic oxidant stress in PPN cholinergic neurons (**Fig. 4h,i**). Taken together, these results suggest that PFFs induce a qualitatively similar set of deficits in dopaminergic and cholinergic neurons at-risk in PD.

**Fig. 4.**
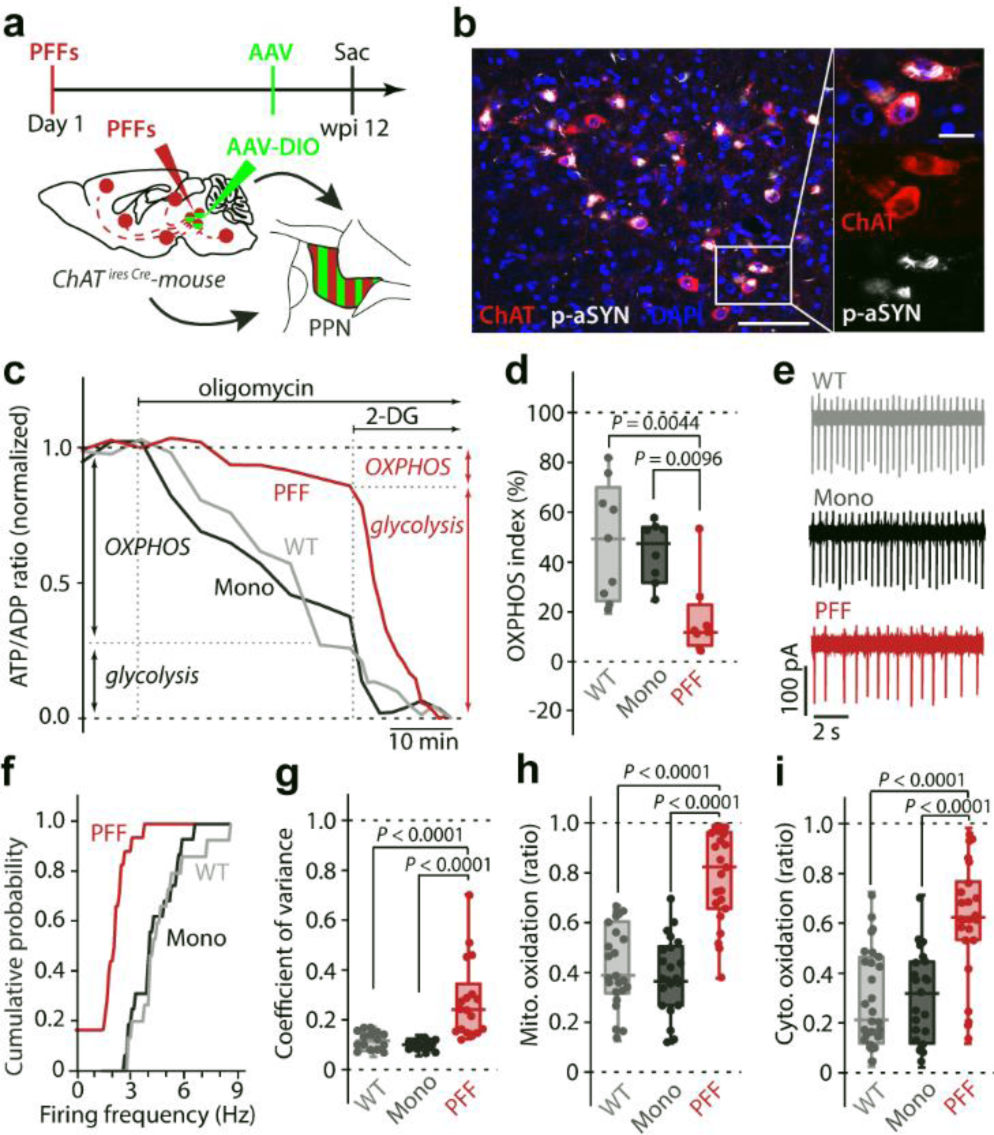
α-synucleinopathy induces similar metabolic, electrophysiological, and oxidative alterations in cholinergic PPN neurons. a, Experimental protocol. **b,** Image depicting p-aSYN pathology in ChAT^+^ PPN neurons 12 weeks after initial seeding. Scale bar, 100 µm in overview, 20 µm in magnified image. **c,** Representative time-lapse measurements of PercevalHR fluorescence ratio. Oligomycin and 2-deoxyglucose (2-DG) were applied to determine OXPHOS and glycolytic contribution to ATP/ADP ratio. **d,** Box plot showing OXPHOS index for WT, Mono, and PFF treated mice. Synucleinopathy shifts neuronal metabolism to the glycolytic pathway (n = 9/8/8 neurons from 6/7/7 mice for PFF, monomeric aSYN, and WT; Kruskal-Wallis test, median ± min/max). **e,** Representative cell-attached recordings of identified ChAT+ PPN neurons. **f,** Cumulative probability plot of ChAT+ PPN autonomous discharge rates (n = 18/16/15 neurons from 6/5/6 mice for PFF, monomeric aSYN, and WT; Kruskal-Wallis, median ± min/max). **g,** Box plot showing coefficient of variance (n = 18/16/15 neurons from 6/5/6 mice for PFF, monomeric aSYN, and WT; Kruskal-Wallis, median ± min/max). **h,** Basal mitochondrial oxidative stress is elevated in cholinergic PPN neurons of PFF injected mice (n = 23/22/23 neurons from 6/5/7 mice for PFF, monomeric aSYN, and WT; Kruskal-Wallis; median ± min/max). **i,** Basal cytosolic ROS levels are significantly increased in cholinergic PPN neurons of PFF injected mice (n = 24/23/28 neurons from 5/5/5 mice for PFF, monomeric aSYN, and WT; Kruskal-Wallis; median ± min/max).

## Discussion

Our studies provide new insight into the cascade of events triggered by synucleinopathy that lead to neuronal dysfunction and degeneration. Twelve weeks after stereotaxic injection of mouse aSYN PFFs in the SNc, roughly a third of the dopaminergic neurons near the injection site had degenerated, and nearly half of the surviving dopaminergic neurons had clearly discernible intracellular aggregates containing PK-resistant pS129 aSYN, as well as p62 – all hallmarks of Lewy pathology [37, 40]. This pathology was limited to PD-vulnerable dopaminergic neurons in the region, leaving neighboring neurons ostensibly unaffected. The appearance of intracellular aSYN pathology was accompanied by a profound disruption in mitochondrial function, as well as clear signs of lysosomal stress. Similar changes were seen in vulnerable PPN cholinergic neurons following PFF seeding.

### aSYN PFF seeding induced cell-specific pathology

In PD, SNc dopaminergic neurons are selectively vulnerable to Lewy pathology and degeneration.

Neighboring neurons in the substantia nigra pars reticulata (SNr), zona incerta and red nucleus are devoid of Lewy pathology [11]. This selective pattern of pathology was recapitulated three months after stereotaxic injection of aSYN PFFs into the SNc. At this time, roughly a third of SNc dopaminergic neurons had been lost. Interestingly, cell counts based upon TH and NeuN expression yielded very similar numbers, suggesting that PFFs did not trigger senescence in a substantial population of SNc dopaminergic neurons [50]. In the surviving dopaminergic neurons, just less than half had perinuclear PK-resistant inclusions that were immunoreactive for pS129 aSYN and p62. This pathology was found throughout the rostrocaudal and mediolateral extent of the SNc and in the VTA, but rarely in neighboring structures. A similar degree of selectivity was found following aSYN PFF injection into the PPN, where PPN cholinergic neurons manifested pS129 aSYN pathology, but not neighboring glutamatergic or GABAergic neurons [6].

Why there was cellular specificity in the pathology induced by aSYN PFFs is unclear. A variety of mechanisms have been proposed to mediate uptake of aSYN fibrils from the extracellular space [51]. But essentially all of these mechanisms, like macropinocytosis, are ones that are common to most (if not all) cell types. Recent work has pointed to the importance of intracellular Ca^2+^ signaling in macropinocytosis and the uptake of aSYN PFF [52, 53]. Indeed, robust intracellular Ca^2+^ signaling and weak intrinsic Ca^2+^ buffering are features that distinguish SNc dopaminergic neurons from neighboring SNr GABAergic neurons [54–56]. Heterogeneity in Ca^2+^ signaling and linked endocytic processes also might be responsible for the apparent resistance of many SNc dopaminergic neurons to aSYN PFF seeding [57].

However, it is important to note that phosphorylation of aSYN at S129 is a relatively late event in the cascade leading to intracellular inclusions [38]. As a consequence, it could be that mesencephalic neurons indiscriminately take up aSYN PFFs but some nominally resistant neurons dispose of them before they escape into the cytoplasm, aggregate and are phosphorylated. It is also possible that the selective distribution of pathology seen in our experiments was dependent upon the timing of the assay (12 wks post injection) and the species of aSYN PFFs used [30, 58]. Nevertheless, the selectivity of the observed pathology argues that cell autonomous factors contribute in a significant way to the distribution of aSYN pathology observed in PD.

### aSYN PFFs induced a complex intracellular pathology

The intracellular pathology induced in dopaminergic neurons by PFFs was varied in composition and structure. EM examination of retrogradely labeled dopaminergic neurons revealed shrunken somata and an increase in the proportion of the cytoplasm occupied by endolysosomes, autophagosomes, and multilamellar terminal lysosomes. Often, these inclusions had fragments of mitochondria, like some types of Lewy pathology [59]. As one might expect, there was a concomitant up-regulation in the expression of genes associated with lysosomal function in dopaminergic neurons as assessed by using the RiboTag method to isolate cell-specific mRNAs [15]. Although there was an alignment with the lysosomal morphology, there are caveats to the transcriptomic studies. One is that the isolation of tagged ribosomes is not perfect, leading to some degree of contamination from other cell types, like astrocytes and microglia, which clearly respond to aSYN PFF seeding [60]. To limit the impact of this shortcoming on the pathway analysis, only mRNAs with counts above 200 were included. Another consideration is that not all of the dopaminergic neurons manifested aSYN pathology, leading to a mix of mRNAs from ostensibly challenged and unchallenged neurons. As a consequence, our profiling may have under-estimated the magnitude of the transcriptome changes induced by aSYN PFFs. This heterogeneity may also explain some puzzling features of the profiles, like the apparent down-regulation in mRNAs for subunits of the lysosomal V-ATPase. Interestingly, while the expression of lysosomal genes was increased by PFF exposure, those linked to the proteasome were down-regulated, in agreement with previous work [46, 47].

### aSYN PFFs disrupted mitochondrial function

Perhaps the most profound change observed in dopaminergic neurons following aSYN PFF seeding was the disruption of mitochondrial OXPHOS. Based upon PercevalHR measurements, mitochondria were making little if any contribution to maintaining cytosolic ATP/ADP ratio, in contrast to control or monomer-exposed neurons. In fact, in many neurons, cytosolic ATP/ADP ratio rose following inhibition of MCV with oligomycin, indicating that mitochondria were importing ATP to stay polarized. Consistent with this functional assay, PFF exposure down-regulated the expression of a wide array of somatic genes coding for OXPHOS proteins and suppressed autonomous spiking, a consequence attributable to an up-regulation in K^+^ channels. Paralleling these changes in activity, the expression of genes related to dopamine synthesis, sequestration and release were also down- regulated by aSYN PFF seeding.

In several respects, these sequelae were like those induced by targeted genetic disruption of MCI function, which leads to a slow, progressive degeneration of SNc dopaminergic neurons and a parkinsonian phenotype in mice [27]. However, there were several significant differences. One difference was that aSYN PFF seeding ostensibly failed to up-regulate the expression of genes coding for proteins involved in glycolysis to compensate for the loss of mitochondrial OXPHOS. It is possible that in those dopaminergic neurons with aSYN pathology, there was an up-regulation in glycolytic genes, but this change was obscured by the inclusion of dopaminergic neurons in the RiboTag sample that did not have aSYN pathology. Another possibility is that the aSYN PFF-induced metabolic remodeling in dopaminergic neurons was constrained by the pleiomorphic nature of the synuclein insult. A key node in the signaling pathway mediating metabolic adaptations is the mammalian target of rapamycin complex 1 (mTORC1) [61]. Recent work has revealed that aSYN fibrils stimulate aberrant mTORC1 activity by binding to tuberous sclerosis protein 2 (TSC2) [62]. In other neurons, mTORC1 up-regulates glycolysis following disruption of mitochondrial electron transport function [63]. Thus, elevated mTORC1 activity could be responsible for the enhanced glycolytic activity in PFF-seeded dopaminergic neurons, independently of any alteration in gene transcription. This metabolic shift also might redirect glycolytic flux away from the pentose phosphate pathway and nicotinamide adenine dinucleotide phosphate (NADPH) generation [64]– compromising cytosolic oxidant defenses and inducing the elevation in cytosolic oxidant stress seen in dopaminergic neurons following aSYN PFF seeding.

Another unresolved question is the nature of the interaction between cytosolic aSYN fibrils and mitochondria. That is, how do aSYN fibrils bring about a deficit in mitochondrial OXPHOS? The physical dimensions of aSYN fibrils precludes the possibility that they are entering mitochondria through outer membrane pores or transporters [59]. Although aSYN fibrils can disrupt mitochondrial membranes in some circumstances [25, 65], this should trigger cytochrome C release, apoptosis and rapid cell death, which is not what appears to be happening in most dopaminergic neurons *in vivo* following aSYN PFF seeding. Another possibility is that the effect of cytosolic aSYN fibrils on mitochondria is secondary to lysosomal or autophagic dysfunction. Lysosomal pathology was a prominent feature of dopaminergic neurons following seeding. It is reasonable to assume that the transit of fibrils from the endolysosomal space into the cytosol reflects lysosomal leakage (at least in part) [60]. As the lysosomal compartment is enriched in Ca^2+^, iron, and proteolytic enzymes, the loss of lysosomal integrity could have direct, deleterious consequences on mitochondria, beyond those mediated by perturbation in mTOR activity and mitochondrial quality control [66]. What is less clear is why lysosomal leakage would be peculiar to dopaminergic or cholinergic neurons. A contributing factor could be the pre-existing burden on the autophagic network in these neurons created by a large axonal arbor and high basal rates of mitochondrial turnover [28, 56, 67].

## Conclusions

Mitochondrial dysfunction and aSYN-enriched LP are hallmarks of PD. But, whether the two are mechanistically linked in a disease relevant setting has been unclear. The studies presented here show that *in vivo* exposure to aSYN PFFs leads to profound, complex intracellular pathology that includes mitochondrial dysfunction in two types of neuron at risk in PD – SNc dopaminergic neurons and PPN cholinergic neurons. These studies point to a pivotal role for mitochondrial dysfunction in aSYN-induced neuronal dysfunction and degeneration, underscoring the potential therapeutic benefit of boosting mitochondrial function in early-stage PD patients [10].

## List of abbreviations

2-DG: 2-deoxyglucose
2PLSM: two photon laser scanning microscopy
AAV: adeno-associated virus
aCSF: artificial cerebrospinal fluid
ADP: adenosine diphosphate
aSYN: alpha-synuclein
ATP: adenosine triphosphate
ChAT: choline acetyltransferase
Cre: Cre recombinase
Cyto: roGFP roGFP targeted to cytosol DAT dopamine transporter
DIO: double-floxed inverse open reading frame
DNA: deoxyribonucleic acid
ETC: electron transport chain
GABAA: gamma-aminobutyric acid type A GSEA gene set enrichment analysis
ISR: induced stress response
MCI: mitochondrial complex I
MCV: mitochondrial complex V
mito-roGFP roGFP: targeted to mitochondrial matrix
mRNAs: messenger ribonucleic acids
NADH: nicotinamide adenine dinucleotide + hydrogen
OXPHOS: oxidative phosphorylation PBS phosphate-buffered saline
PD: Parkinson’s disease
PFA: paraformaldehyde
PFFs: pre-formed alpha-synuclein fibrils
PINK1: PTEN induced kinase 1
PK: proteinase K
PPN: pedunculopontine nucleus
pS129: phosphorylation at serine 129
roGFP: redox-sensitive variant of green fluorescent protein
SNc: substantia nigra pars compacta
SNr: substantia nigra pars reticulata
TEM: transmission electron microscopy
TH: tyrosine hydroxylase
ThT: thioflavin T
WT: wildtype

## Declarations

### Ethics approval

All animal experiments were performed according to the NIH Guide for the Care and Use of Laboratory Animals and approved by the Northwestern University Animal Care and Use Committee.

### Consent for publication

Not applicable

### Availability of data and materials

The datasets generated and/or analyzed during the current study are available in the Zenodo repository (DOI: 10.5281/zenodo.10251490).

### Competing interests

The authors declare that they have no competing interests.

### Funding

This study was supported by awards to DJS by the JPB Foundation, NIH (NS 121174) and Aligning Science Across Parkinson’s [ASAP-020551] through the Michael J. Fox Foundation for Parkinson’s Research (MJFF). TMD was funded by Aligning Science Across Parkinson’s [ASAP-020608] through the Michael J. Fox Foundation for Parkinson’s Research (MJFF). FFG received a grant from StichtingParkinson Fonds and the Von-Behring-Röntgen Stiftung. WHO is supported by the Charitable Hertie Foundation, Frankfurt/Main, Germany.

### Authors’ contributions

DJS, MTH, FFG and WHO designed the experiments, wrote, and edited the manuscript. FFG, and MTH performed experiments, conducted data analysis, and created the figures. EZ and DW assisted with the imaging experiments. ZX and TK assisted with the molecular profiling.

CG and EN performed histological analysis and electron microscopy. VLD and TMD provided PFFs and guidance on their handling and use. ND provided guidance on the analysis and interpretation of the RNAseq experiment. All authors have read and approved the manuscript.

## Acknowledgements

We wish to thank Sasha Ulrich and Sabine Anfimov for expert technical assistance, Katrin Roth and the Core Facility Cellular Imaging of Philipps University-Marburg for the use and assistance with the Leica TCS SP8 confocal microscope, and Frederik Helmprobst and the Core Facility for Mouse Pathology and Electron Microscopy for assistance with TEM experiments.

---

**Suppl. Fig. S1.**
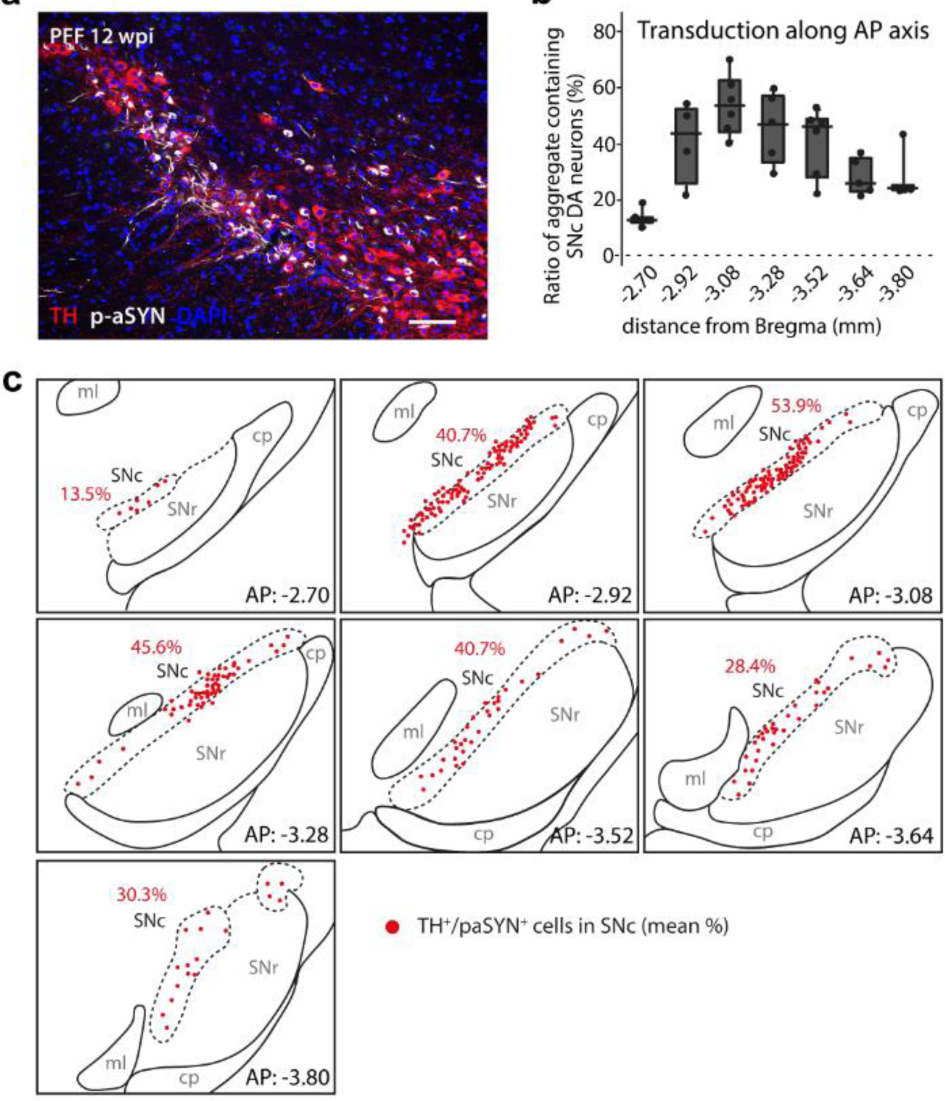
Homogenous distribution of seeded α-synucleinopathy in DA SNc of DAT-Cre-WT mice. a, TH^+^ SNc neurons exhibiting p-aSYN pathology 12 weeks after initial seeding. **b,** Boxplot showing distribution of TH^+^ SNc neurons harboring p-aSYN aggregates over the rostro-caudal extent of the SNc (n = 3/4/6/5/6/5/3 sections from 7 mice; median ± min/max). **c,** Percentage of paSYN^+^ TH^+^ cells depicted for the analyzed AP-coordinates. Abbreviations: cp, cerebral peduncle; ml, medial lemniscus; SNc, substantia nigra pars compacta; SNr, Substantia nigra pars reticulata.

**Suppl. Fig. S2.**
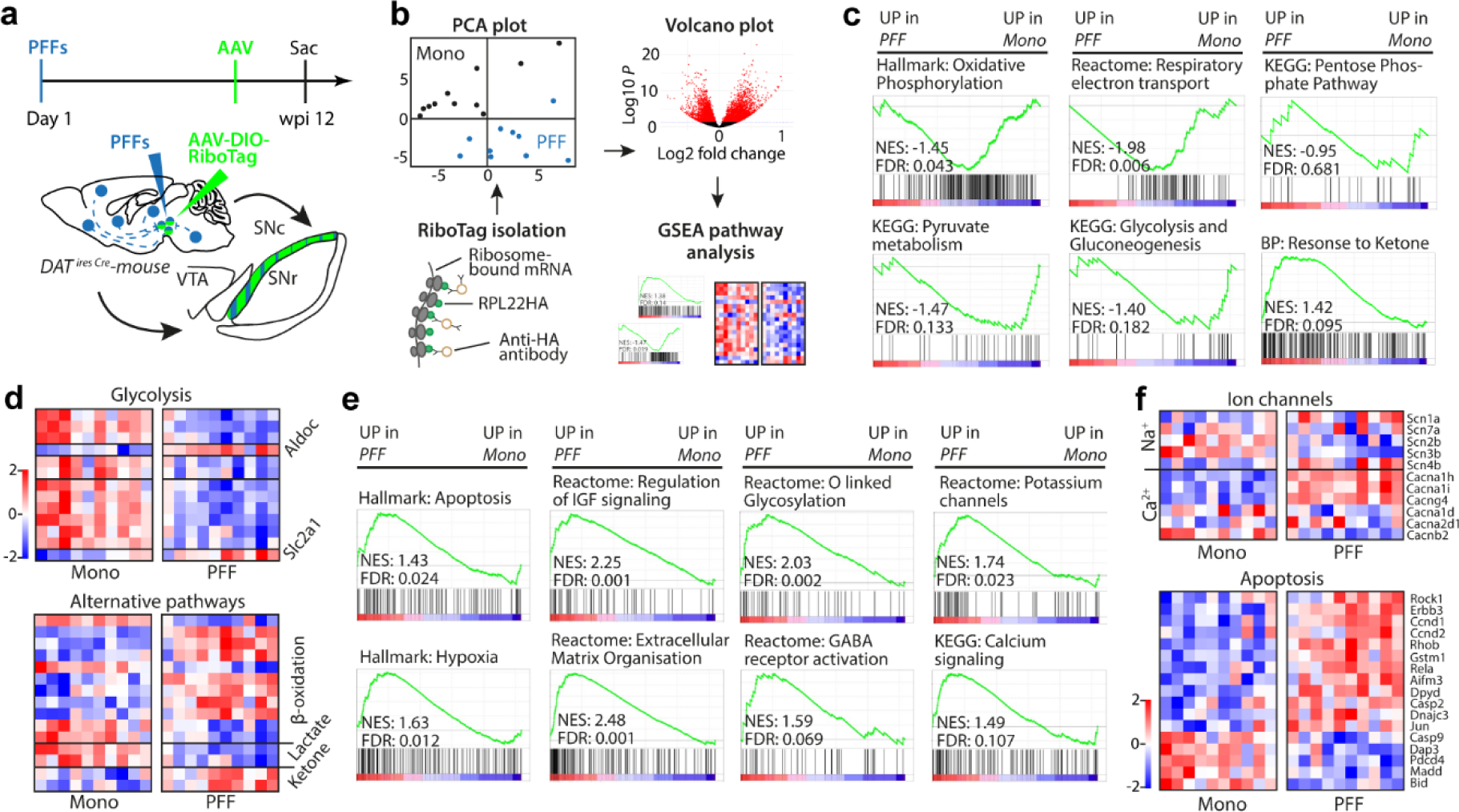
RNAseq workflow and pathway analysis using GSEA. a, Experimental protocol. **b,** RNAseq workflow including RiboTag isolation, quality control including PCA plot, analysis of differential gene expression and pathway analysis using GSEA. **c,** Gene set enrichment analysis (GSEA) for metabolic related pathways in DA SNc neurons from mice injected with either aSYN PFF or monomeric aSYN (n = 10 mice for aSYN PFF or monomeric aSYN injected DAT-Cre-WT mice). **d,** Heatmaps of RNASeq analysis showing significantly down- or upregulated genes of glycolysis, *β*-oxidation, lactate metabolism, and ketone body consumption (n = 10 for both groups; Wald test adjusted using Benjamini-Hochberg method, p < 0.05). **e,** Plots depicting GSEA for highly enriched pathways in aSYN PFF injected group (n = 10 mice for aSYN PFF or monomeric aSYN injected DAT-Cre-WT mice). **f,** Heatmaps of RNASeq analysis showing significantly down- or upregulated genes of certain ion channels and apoptosis (n = 10 for both groups; Wald test adjusted using Benjamini-Hochberg method, p < 0.05). Abbreviations: KEGG, Kyoto Encyclopedia of Genes and Genomes; FDR, false discovery rate. NES, normalized enrichment score; y-axis, enrichment score; x-axis, rank in ordered dataset.

**Suppl. Fig. S3.**
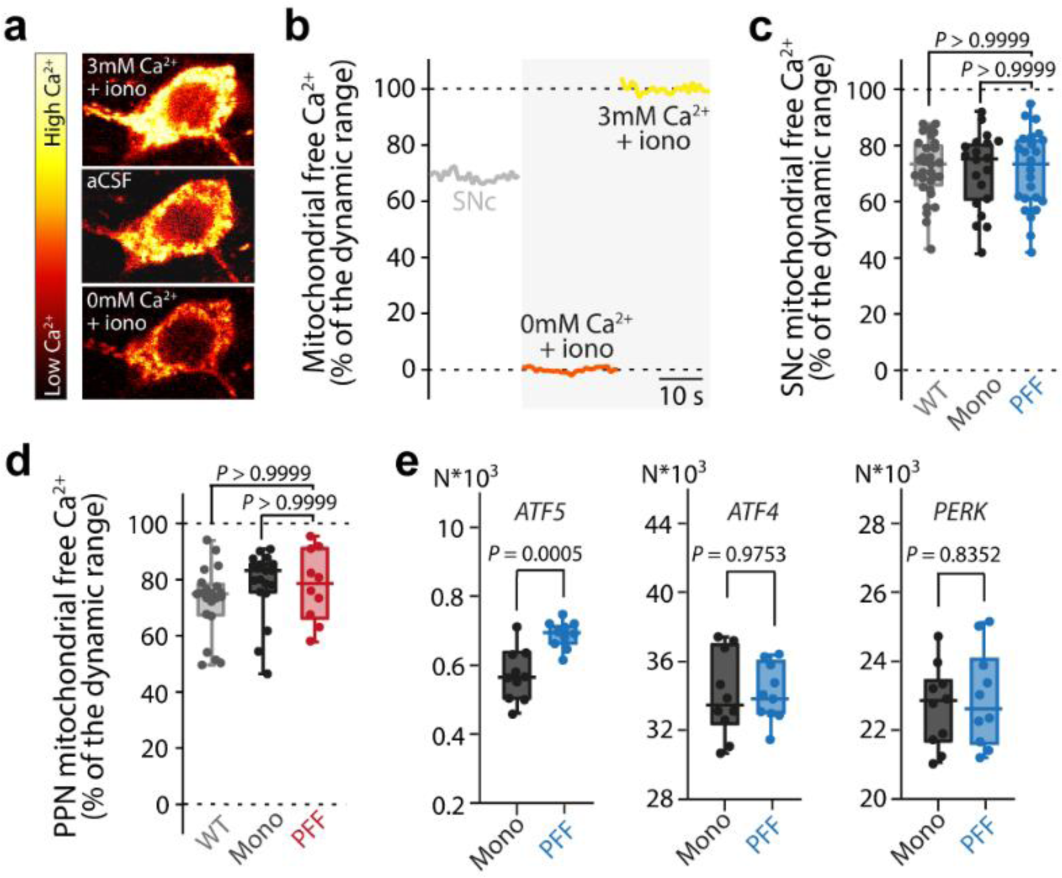
Mitochondrial Ca2+-levels and Induced Stress Response (ISR) genes. a,b, The dynamic range of the expressed mito-GCaMP6 probe was determined by the addition of the Ca^2+^-ionophore ionomycin (iono) in aCSF with 0 mM Ca^2+^ (lowest Ca^2+^-level) followed by the application of aCSF with 3 mM Ca^2+^ (highest Ca^2+^-level). **c,** Mitochondrial Ca^2+^-levels in DA SNc neurons were unaffected by induced synucleinopathy (n = 24/20/31 from 6/5/5 mice for aSYN PFF, monomeric aSYN, and WT; Kruskal-Wallis; median ± min/max). **d,** Mitochondrial Ca^2+^-levels in CN PPN neurons were unaffected by induced synucleinopathy (n = 10/19/21 from 5/5/5 mice for aSYN PFF, monomeric aSYN, and WT; Kruskal-Wallis; median ± min/max). **e,** Box plots depicting normalized gene expression values for ISR genes (ATF5, ATF4, PERK) highlighting upregulation of transcription factor ATF5 (n = 10 for both groups; Wald test adjusted using Benjamini-Hochberg method, median ± min/max).

**Suppl. Fig. S4.**
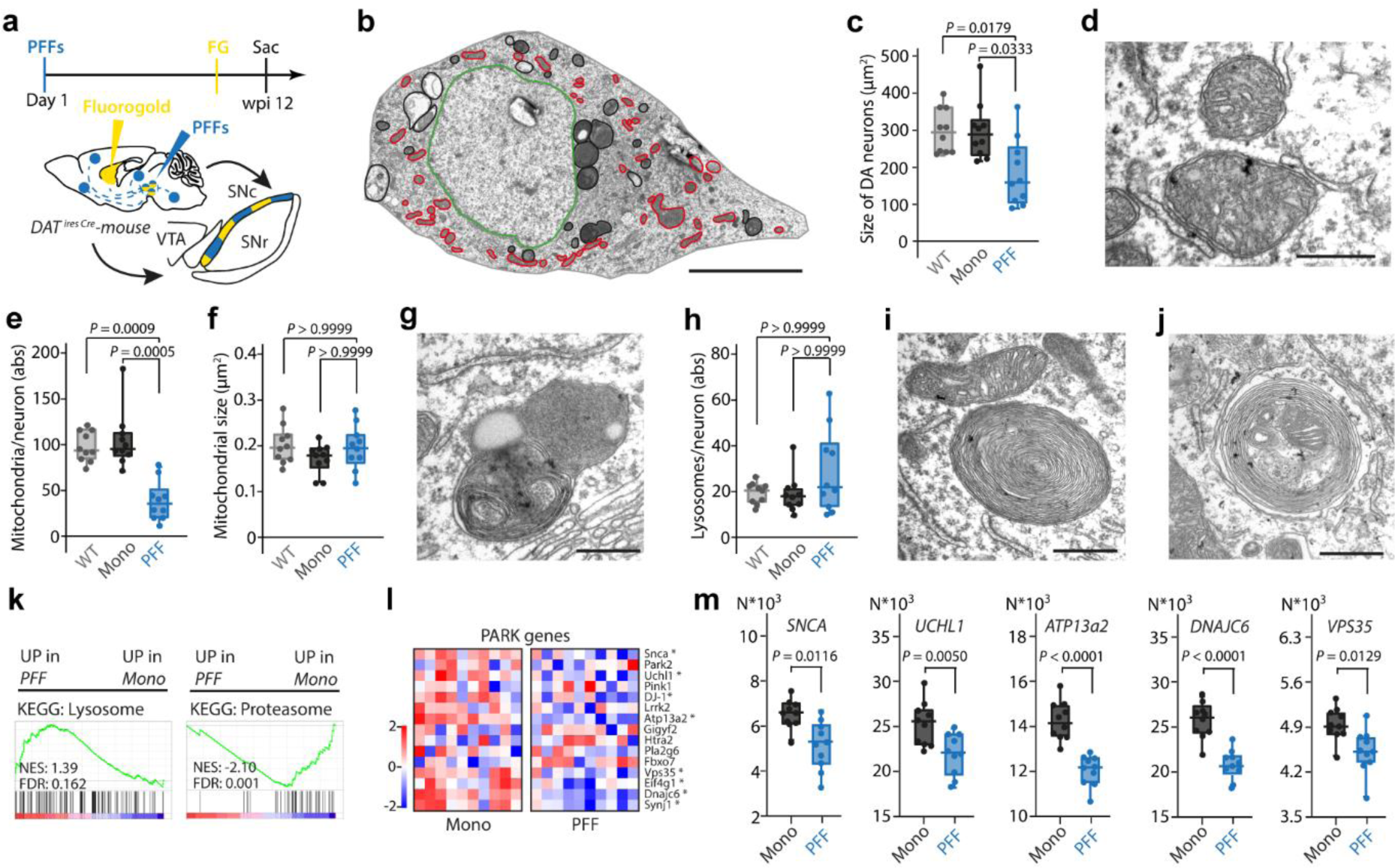
aSYN PFF induced α-synucleinopathy causes morphological alterations of mitochondria and lysosomes. a, Experimental protocol. **b,** Transmission electron micrograph (TEM) of a SNc DA neuron from an aSYN PFF injected mouse, showing large perinuclear pathology. The nucleus is highlighted in green, mitochondria in red, and lysosomes in black, respectively. Scale bar 10 µm. **c,** Quantification of DA SNc neuronal soma size (n = 10/10/10 neurons from 4/4/4 mice for aSYN PFF injected DAT-Cre-WT, and monomeric aSYN injected DAT-Cre-WT mice, and C57Bl6/J-WT mice; Kruskal-Wallis; median ± min/max). **d,** Dysmorphic mitochondria within a DA SNc neuron from an aSYN PFF injected DAT-Cre-WT mouse. Scale bar 500 nm. **e,f,** Quantification of mitochondria numbers (**e**) and size of mitochondria per SNc DA neuron (n = 10/10/10 neurons from 4/4/4 mice for aSYN PFF injected DAT-Cre-WT, and monomeric aSYN injected DAT-Cre-WT mice, and C57Bl6/J-WT mice; Kruskal-Wallis; median ± min/max). **g,** TEM depicting activated lysosome within a DA SNc neuron from an aSYN PFF injected DAT-Cre-WT mouse. Scale bar 500 nm. **h,** Quantification of lysosomes per DA SNc neuron n = 10/10/10 neurons from 4/4/4 mice for aSYN PFF injected DAT-Cre-WT, and monomeric aSYN injected DAT-Cre-WT mice, and C57Bl6/J-WT mice; Kruskal-Wallis; median ± min/max). **i, j,** TEM images from SNc DA neurons from an aSYN PFF injected mice depicting a lamellar accompanied by a mitochondrion with swollen cristae structure (**i**), and a lamellar body with a trapped mitochondrion. Scale bars 500 nm. **k,** Gene set enrichment analysis (GSEA) of DA SNc neurons from DAT-Cre mice injected with either aSYN PFF or monomeric aSYN. KEGG pathway “Lysosome” was upregulated in aSYN PFF injected mice, while KEGG pathway “Proteasome” was strongly downregulated (n = 10 mice for aSYN PFF or monomeric aSYN injected DAT-Cre-WT mice). KEGG, Kyoto Encyclopedia of Genes and Genomes; FDR, false discovery rate. NES, normalized enrichment score; y-axis, enrichment score; x-axis, rank in ordered dataset. **l,** Heatmap of RNASeq analysis showing expression of PARK genes (n = 10 for both groups; Wald test adjusted using Benjamini-Hochberg method, p < 0.05). **m,** Box plots showing normalized gene expression values for significantly changed PARK genes (SNCA, UCHL1, ATP13a2, DNAJC6, VPS35). Note, all genes, except for SNCA, are implicated in lysosomal or proteasomal degradation of proteins (n = 10 for both groups; Wald test adjusted using Benjamini-Hochberg method, p < 0.05; median ± min/max).

**Suppl. Fig. S5.**
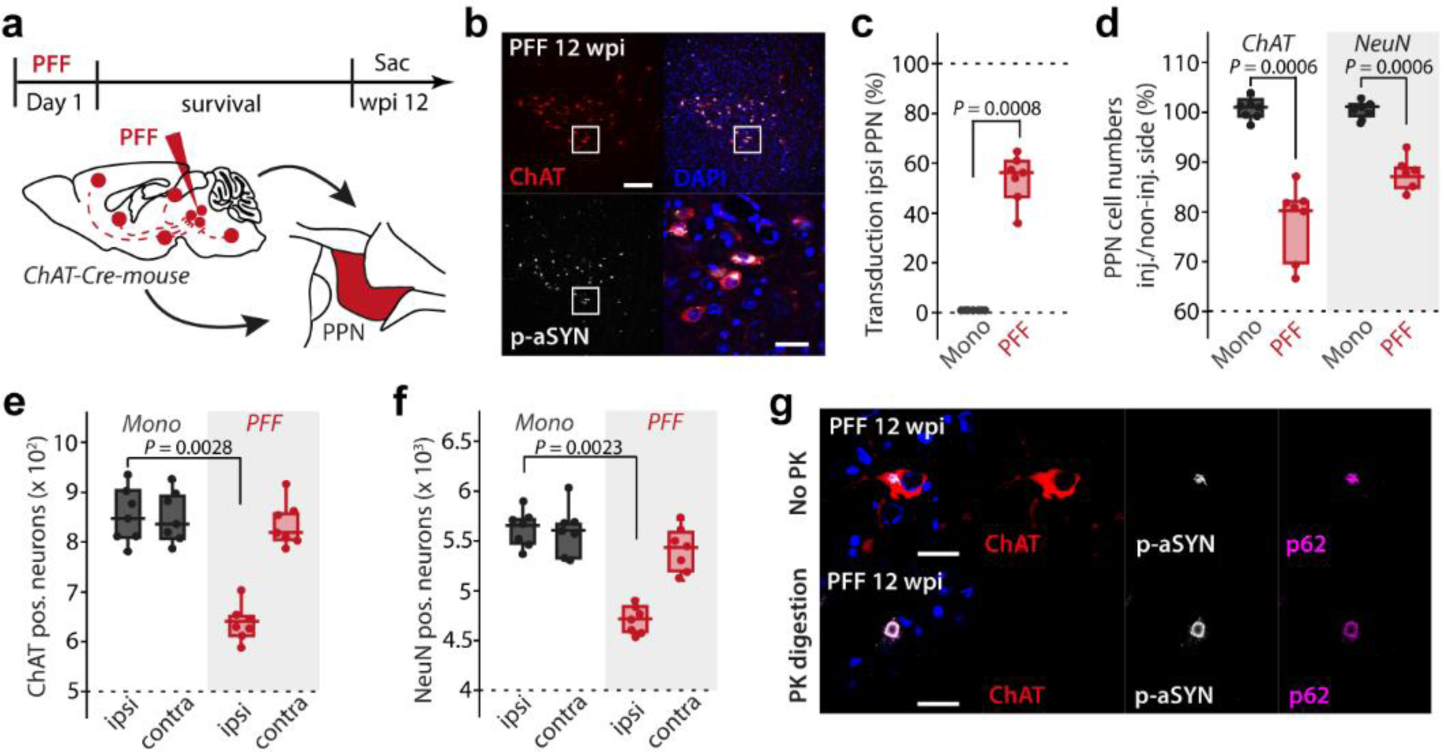
aSYN PFF injection causes PD-like neurodegeneration of PPN cholinergic neurons. a, Experimental protocol. **b,** ChAT+ PPN neurons exhibiting p-aSYN pathology 12 weeks after initial seeding. Scale bar, 250 µm in overview, 25 µm in magnified image. **c,** Transduction rate for PPN injected mice on the ipsilateral side of injection (n = 7 for both groups; Mann-Whitney-U; median ± min/max). **d,** Quantification of ChAT+ and NeuN+ PPN cells expressed as percentage of the non-injected side (n = 7 for both groups; Mann-Whitney-U; median ± min/max). **e,** Box plot showing number of ChAT+ neurons in PPN (n = 7 for both groups; Kruskal-Wallis; median ± min/max). **f,** Box plot showing number of NeuN+ neurons in PPN (n = 7 for both groups; Kruskal-Wallis; median ± min/max). **g,** p-aSYN+ aggregates in ChAT+ PPN neurons were p62+ and resistant to digestion with Proteinase K (PK). Scale bar, 25 µm.

## Notes

### Competing Interest Statement

The authors have declared no competing interest.

